# A Closed-Form Bayesian Model for DNA Replication Reveals Intrinsic Origin Timing and Activation Delays

**DOI:** 10.64898/2026.05.28.728365

**Authors:** Dario D’Asaro, Diletta Ciardo, Laurent Lacroix, Alan Tatourancheau, Bertrand Theulot, Olivier Hyrien, Benoît Le Tallec, Arach Goldar, Benjamin Audit, Jean-Michel Arbona

## Abstract

We present an analytical framework for modeling eukaryotic DNA replication that, given experimental Replication Fork Directionality (RFD) data, enables Bayesian inference of origin number, activation delay 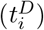 and intrinsic timing (*λ*_*i*_), (the mean replication time if each origin were isolated). By deriving closed-form expressions for RFD and Mean Replication Timing (MRT) under exponential and a specific Weibull firing-time distributions as functions of 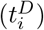 and (*λ*_*i*_), we eliminate the need for stochastic simulations.

These analytical results reveal that RFD, as a ratio of fork directions, is invariant under joint rescaling of intrinsic timing and fork speed; absolute intrinsic timing can nonetheless be inferred when fork speed is independently measured.

We demonstrate that under exponential firing distribution for the origin, the observed efficiency (*E*_*i*_), i.e. the probability for an origin to fire which accounts for nearby origin, is simply MRT(*x*_*i*_)*/λ*_*i*_.

The closed-form RFD expressions allow to use a Bayesian method that achieves 0.96–0.99 correlation with yeast RFD profiles and resolves ~780 origins in *S. cerevisiae*.

Our framework identifies about 150 origins with biologically significant delays (≥ 3 minute), revealing regulated activation kinetics undetectable by existing methods. By quantifying how origin intrinsic timing and delays shape replication timing landscapes, this work confirms yeast as a paradigm organism for studying DNA replication control mechanisms.

## II. INTRODUCTION

### Challenges of Eukaryotic DNA Replication

DNA replication is a fundamental process ensuring genomic stability during cell division. In eukaryotes, DNA replication initiates at hundreds to thousands of origins, with tightly regulated timing and efficiency [1].

Two complementary experimental approaches provide insights into replication dynamics. The first comprises ChIP-seq assays that measure the chromatin binding of replication initiation complexes, such as the Origin Recognition Complex (ORC) and the Minichromosome Maintenance (MCM) helicase [2–6]. These factors act during G1 phase, when ORC, Cdc6, and Cdt1 load the MCM helicase onto DNA, licensing origins and establishing the pool of potential firing sites. Their genomic occupancy thus serves as an experimental proxy for an origin’s intrinsic timing *λ*_*i*_ [2], which describes how quickly an origin would fire in the absence of neighbouring origins. Importantly, intrinsic timing is not determined solely by factor binding: it can be modulated by chromatin regulators such as Sir2, which alter the local protein state and thereby affect licensing efficiency [7, 8]. The second category comprises sequencing-based assays such as OK-Seq [9] and FORK-Seq [10], which measure population-level quantities including Replication Fork Directionality (RFD) and Mean Replication Timing (MRT). Unlike ChIP-seq, these readouts capture the *net outcome* of replication, reflecting intrinsic origin properties as modulated by neighborhood effects.

During S-phase, licensed origins may be silenced by incoming replication forks through the passivation mechanism: if one origin fires first, the advancing fork can replicate and thereby passivate its neighbors before activation occurs. This phenomenon gives rise to the concept of observed efficiency *E*_*i*_, the probability for an origin to fire accounting for neighboring origins throughout S-phase. The central modelling challenge is thus to connect the intrinsic timing *λ*_*i*_ (accessible via ChIP-seq proxies) to the population-level observables MRT and RFD, disentangling intrinsic origin properties from the neighborhood effects captured by passivation.

### Advances in Experimental Techniques

Recent experimental breakthroughs, such as OK-Seq [9] and FORK-Seq [10], have enabled genome-wide mapping of Replication Fork Directionality (RFD), initiation density [11], and fork speeds [12] in asynchronous cell populations. These methods bypass synchronization-induced perturbations, offering unprecedented insights. Single-molecule studies by Gauthier et al. [13] further demonstrated how local origin firing patterns influence replication dynamics. Despite these advances, computational methods to disentangle the intrinsic origin timing along the genome (*λ*_*i*_) with its observed efficiency (*E*_*i*_) remain limited.

### Limitations of Existing Computational Approaches

Prior computational approaches face limitations. For example, Bazarova et al. [14] used stochastic simulations based on Retkute et al.’s model [15] to estimate origin efficiencies from RFD data. However, their reliance on simulating thousands of cells restricted analyses to small systems of three origins at a time, neglecting interdependencies between neighboring origins. Similarly, Arbona et al. [16] combined neural networks with replication timing data but required computationally intensive iterative training. Bazarova et al. [14] also showed using Bayesian methods that reduced licensing probabilities and origin passivation contribute to replication control, yet these approaches remain constrained by computational bottlenecks.

A recent study by Berners-Lee et al. [17] advanced the field by developing a stochastic, whole-genome model of *S. cerevisiae* replication timing in which origins compete for a limited pool of firing factors. At kilobase resolution, their model successfully reproduced mean replication timing, inter-origin distances, origin efficiencies, and replication fork directionality. This work strongly supports the view that competition among origins, together with limiting firing factors, is a major determinant of replication timing. Nevertheless, the model relies on extensive stochastic simulations, limiting its ability to provide closed-form links between intrinsic strength, observed efficiency, and experimental readouts such as MRT and RFD. Berkeimer et al. [18] recently developed and analytical model for MRT in an exponential firing context, and where the fit was performed using an iterative refinement scheme.

### Our Contributions

We address the limitations of stochastic simulation by developing an analytical framework. It provides closed-form solutions for RFD and MRT, the observed effiency and the standard error on the MRT, under both exponential and more biologically realistic Weibull (with parameter k=2) firing-time distributions. It also allows accounting for potential activation delay due to regulation mechanism 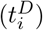. This approach eliminates the computational expense of stochastic simulations, enabling rapid, genome-wide Bayesian inference of intrinsic origin timing (*λ*_*i*_) and activation delays. Our model yields critical mechanistic insights, demonstrating that RFD discontinuities directly encode the observed, passivation-adjusted firing efficiency (*E*_*i*_). We establish the theoretical relationship between RFD, *λ*_*i*_, and fork speed (*v*), allowing for the inference of absolute origin timing when fork speed is known. Furthermore, under an exponential firing model, we derive the simple but powerful relationship *E*_*i*_ = MRT(*x*_*i*_)*/λ*_*i*_, which analytically clarifies the often-conflated distinction between intrinsic timing and observed efficiency as inferred from RFD data. Applying this scalable framework to *S. cerevisiae*, we resolve approximately 780 origins and identify about 145 with biologically significant activation delays (≥ 3 minute), revealing a layer of regulated activation kinetics previously undetectable by other methods.

### Significance

Our work bridges the gap between theoretical models and data-driven analysis, offering a robust tool to study replication dynamics under diverse conditions. By distinguishing **intrinsic origin timing (***λ*_*i*_) from **observed efficiencies (***E*_*i*_), with a simple relationship, we clarify a critical distinction. We validate our framework on synthetic data and apply it to experimental yeast datasets, revealing biologically significant delays in about 150 origins. This framework establishes yeast as a paradigm system for dissecting DNA replication control mechanisms. Studies of genetic variation in replication timing underscore the importance of such mechanistic models for understanding genomic stability [19].

## III. MODEL

Retkute et al. [15] model replication using discrete origins along a chromosome. The probability that origin *i* is activated is *q*_*i*_*p*(*t*, Λ_*i*_), where *p*(*t*, Λ_*i*_) is a probability distribution parameterized by Λ_*i*_, and *q*_*i*_ describes the licensing efficiency of the origin. While in the following sections we set the licensing probability to 1, we will address the effects of relaxing this assumption later. Note that even for *q*_*i*_ = 1, an origin may still not fire due to its passivation.

The probability *p*(*t*, Λ_*i*_) represents an arbitrary probability distribution of firing, but later on will be fixed to be exponential or Weibull with parameter *k* = 2. If *t <* 0 then *p*(*t*, Λ_*i*_) = 0. The probability for a fork starting from origin *i* located at position *x*_*i*_ to reach the point *x* at *t* is:

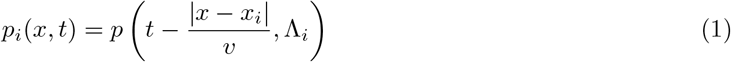

which is a delay of the initial probability distribution due to the propagation of the fork.

The probability that another fork *j* arrives at *x* later than *t* is given by

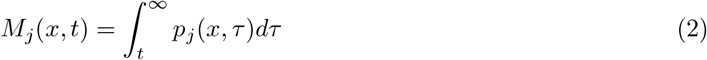

Starting from these equations it can be shown App. A that:

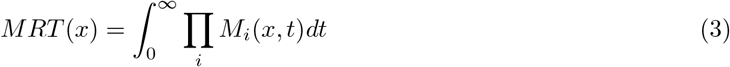

which is considerably simplified with respect to the original formulation of Retkute et al. [15] which combines both Eq. (A6) and Eq. (A4). This allows the analytical derivation of MRT and 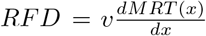 and *E*_*i*_, the observed efficiency of the origin *i* for two types of distribution of origin firing.

We first evaluate these quantities for the exponential distribution, followed by the Weibull case with shape parameter *k* = 2. These two probability distributions are plotted in Fig 1 A,E respectively.

**FIG. 1.**
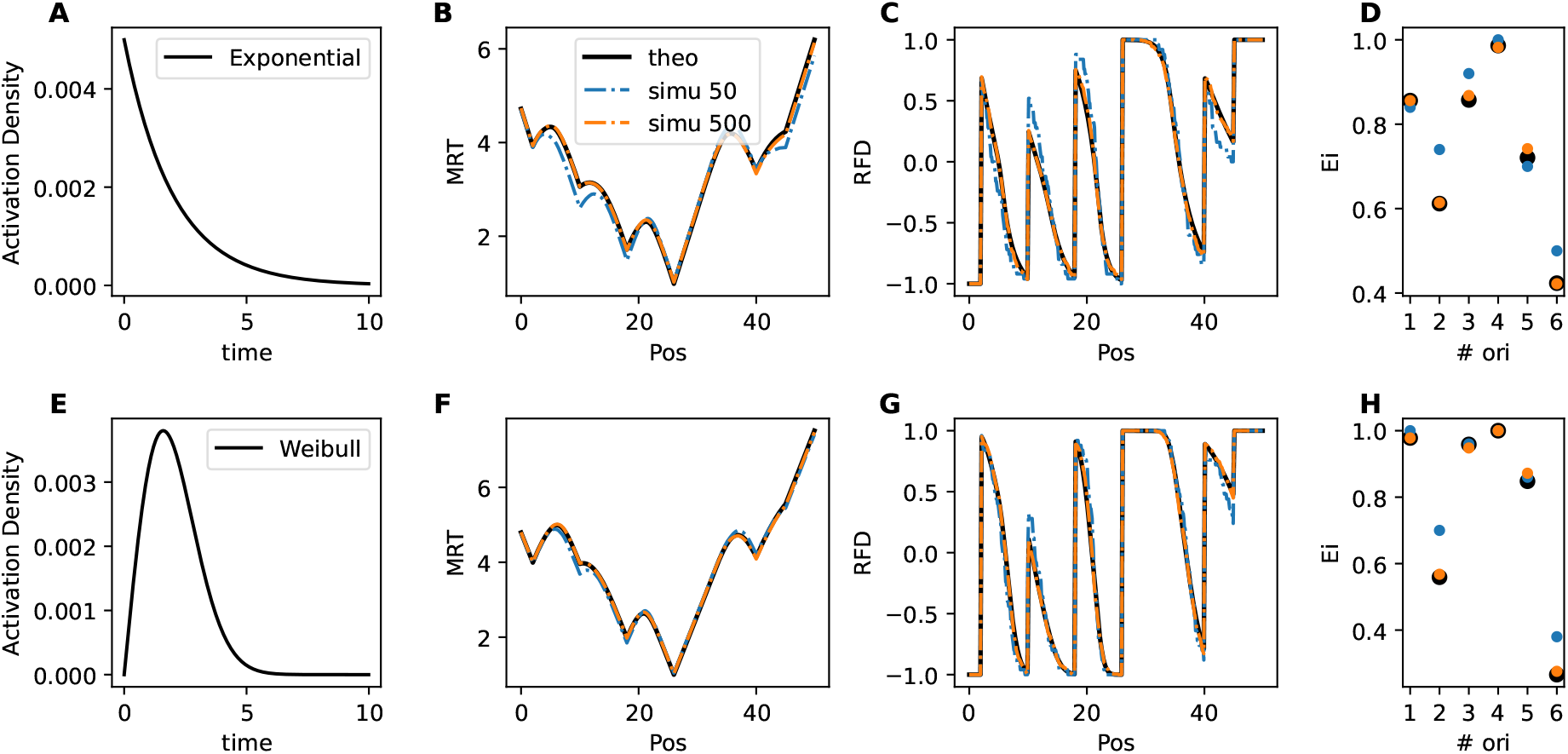
Analytical vs. simulated profiles for exponential (A–D) and Weibull *k* = 2 (E–H) models. **(A/E)** Activation function for *λ* = 2,**(B/F)** MRT, **(C/G)** RFD, and **(D/H)** observed efficiencies *E*_*i*_ show near-perfect agreement. Simulations involve stochastic firing time sampling and fork propagation at constant speed, with origins passivated (silenced) if preempted by neighboring forks. For both models, 50 simulations (blue dotted lines) exhibit residual noise, while 500 simulations (orange dotted lines) converge to the analytical solution (black lines). Here are the chosen parameter values for the comparison : *λ* = [0.50, 5.00, 2.00, 1.00, 3.33, 10.00] (min) *t*^*D*^ = [3.50, 0.00, 0.00, 0.00, 1.00, 0.00] (min), and position of the origins: [2.00, 10.00, 18.00, 26.00, 40.00, 45.00] (kb) and fork speed *v* = 2.5kb/min. The same parameters where chosen for the exponential and Weibull case.

### A. Exponential Firing Distribution

The replication dynamics for a system of *N* origins with exponentially distributed firing times yields closed-form solutions for MRT and RFD. The firing probability for origin *i* is 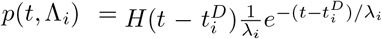, with H(t) the Heaviside function (Fig 1 A). So each origin is parametrized by two values, the scale parameter *λ*_*i*_ and a hard delay of activation 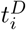.

We define 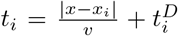 at *x*, as the total delay for a fork from an origin at *x*_*i*_ to reach position *x*.This delay comprises two components: the propagation time 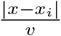 (time for the fork to travel the distance |*x* − *x*_*i*_| at speed *v*) and an optional activation delay 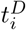 (mimicking regulatory delays in origin firing).

To compute MRT(*x*), we first sort the *n* origins by their earliest arrival times *t*_*i*_ (ascending order). The integration is partitioned into intervals where only subsets of origins contribute. Between *t* ∈ [0, *t*_1_), all *M*_*i*_(*t, x*) = 1 (no forks have arrived), while for *t* ∈ [*t*_*i*_, *t*_*i*+1_), only the first *i* origins contribute non-trivially.

After calculation (App. B), the MRT at position *x* is:

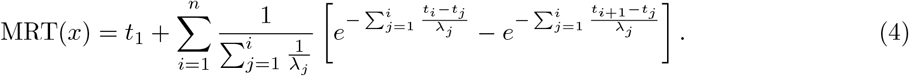

Similarly it is possible to compute the variance of the MRT (App. B).

a. *RFD Derivation:* The RFD is derived as the spatial derivative of MRT scaled by fork speed:

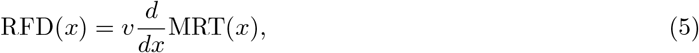

where differentiation accounts for the linear dependence 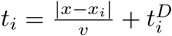 (see App. B for details).
b. *Observed Efficiency (E*_*i*_): The observed efficiency or the passivation-adjusted firing probability is defined as: 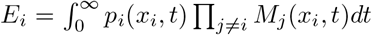 which corresponds to the probability that the fork *i* is activated multiplied by the probabilities that other forks arrive later, and this integrated over the whole S-phase. The exponential model yields (App. B):

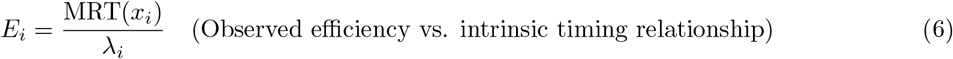

valid when 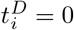. While not apparent for this relationship, *E*_*i*_ is indeed a probability bounded between 0 and 1 as MRT(*x*_*i*_) *< λ*_*i*_. Delays 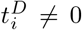 invalidate this simple relationship if forks preempt origin activation (App. B).

### B. Weibull Firing Distribution (shape parameter *k* = 2)

The Weibull distribution, parameterized by a shape parameter *k* and a scale parameter *λ*, allows for gradual origin activation. For *k* = 2, it takes the form 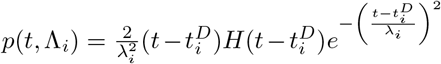 as shown in Fig 1 E. Unlike the sharp transition of the exponential case, this distribution captures a progressive increase in firing at 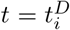, followed by a decrease. It is to be noted that the Weibull distribution with shape parameter *k* = 1 is the Exponential distribution. These two distributions where the only ones that we could resolve analytically.

The Weibull probability distribution (with *k* = 2) also depends on two parameters characterizing the origins: *λ*_*i*_ similar to a scale as in the exponential case, and a hard delay of activation 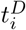. The MRT becomes:

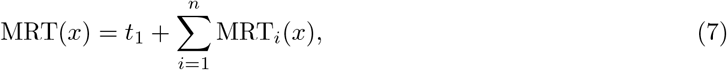

where each term MRT_*i*_(*x*) involves Gaussian integrals:

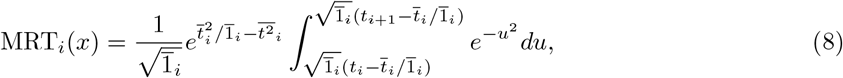

with 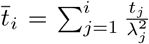 and 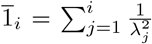. While these integrals involve the error function (erf), they are computationally efficient in practice: standardized approximations for erf are implemented in numerical libraries, eliminating the need for explicit numerical integration during evaluation. The RFD derivation parallels the exponential case but requires differentiating error function terms. (App. C).

Here we used the standard parameterization of the Weibull distribution. However, contrary to the exponential distribution where *λ* is also the mean value of the distribution, for Weibull (*k* = 2)

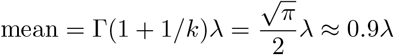

So in practice, we replace *λ* by 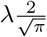, so that the distribution is also parametrised by its mean.

### C. Model validation: comparison with simulations

For computing any quantity at a given point x, it amounts in practice to compute *t*_*i*_ from each origin using parameters *k*_*i*_ and 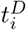 and then computing the various sums. One of the main differences with simulation is that it is possible to compute any quantity at a given point without having to compute the quantities on the full chromosome. In Fig 1 we compare Monte Carlo simulation with N=50/N=500 replicates to the analytical formula. We choose parameters by hand to have combination of origin strength and delay (Fig 1 legend). For the simulation, the activation time of an origin is drawn according to its corresponding activation law. If an activation time is higher than the time it takes for another origin to reach the position of the first mentioned origin, this origin is passivated. We can observe that a simulation with 500 replicates perfectly overlaps with the analytical formula. The difference between the two firing models are small but can be observed as a smoother MRT and RFD profile for the Weibull distribution.

### D. Including partial licensing of origins

To simplify the analytical framework, the model assumes that all origins succeed in licensing. From experimental evidence, it is unclear if that it is the case. Two main questions then arise: (i) can partial licensing be incorporated into the analytical framework, and (ii) if it is present in the data, what is the impact of fitting with a model that does not account for it? Indeed, it is possible to extend the computation to the case *q*_*i*_ *<* 1. The resulting MRT, is then a sum over all 2^*n*^ possible subsets of licensed origins, weighted by their probability:

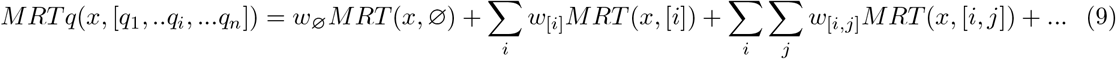

with *MRT* (*x*, ∅), the classical MRT where no origin has been excluded, *MRT* (*x*, [*i*]), the MRT where we discard origin *i* from the set of origin, and so on. The weights are: *w*_∅_ = Π_*j*_ *q*_*j*_, *w*_[*i*]_ = Π_*j*=*i*_ *q*_*j*_(1 *q*_*i*_). The sum over the weights is by construction equal to one.

The full MRT is thus a combinatorial sum of all possible combinations of active origin, whose computation would be impractical. However, if one make the hypothesis that most of the origin have a rather high probability to have licensing, it is possible to approximate the full MRT by adding perturbation due to one origin being not licensed for all origins:

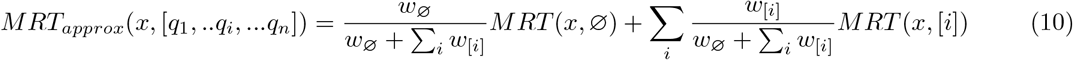

In practice this could be implemented by computing *MRT* (*x*, ∅) and only part of *MRT* (*x*, [*i*]) where the origin *i* was active and should be removed from the computation, the other parts of the MRT being the same. In the following work we did not implement this correction. We will discuss the impact of this hypothesis on the discussion.

## IV. ANALYSIS OF RFD

### A. RFD information content: Inferring Absolute Origin Efficiencies from RFD Data

Our analytical result allows us to reveal an interesting symmetry: RFD profiles under exponential firing are invariant to simultaneous rescaling of fork speed (*v* → *fv*), scale parameters (*λ*_*i*_ → *λ*_*i*_*/f*) and delay 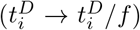 for any *f >* 0. This symmetry arises because the RFD formula includes terms like 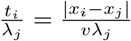, as well as ratio of *λ*_*i*_ which remain unchanged under such transformations. Consequently, RFD alone can resolve both *v*, *λ*_*i*_ and 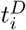 up to a multiplicative constant. However, if the fork speed *v* is independently determined (e.g., experimentally measured), this constant is determined, and RFD encodes *absolute* intrinsic efficiencies through the *passivation mechanism*.

### B. Estimating model parameters

#### 1. Estimation and Model Selection

To infer the model parameters (intrinsic timing *λ*_*i*_ and delays 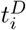) from experimental RFD data, we need a robust statistical method. A straightforward approach is to find the parameters that minimize the difference (e.g., mean squared error) between our model’s predicted RFD profile and the experimental one. While fast, this method does not easily allow us to compare the performance of fundamentally different models, such as the exponential versus the Weibull firing model.

To address this, we adopted a Bayesian framework. This approach not only estimates the likeliest value for each parameter but also quantifies the uncertainty of that estimate. Crucially, it provides a formal way to compare different models. We explored two computational techniques to implement this approach, Automatic Differentiation Variational Inference (ADVI) and the Laplace approximation [20, 21].

[22].

These methods allow us to calculate the Evidence Lower Bound (ELBO), a score that reflects how well a given model explains the experimental data while penalizing overfitting. By comparing ELBO scores, we can select the best model (e.g., Weibull with delays) and the optimal number of origins. While variational methods like ADVI and Laplace can slightly underestimate parameter uncertainty [23], they proved highly effective for model selection in our tests. We found that the Laplace method provided a good balance of speed and accuracy, making it ideal for our genome-wide analysis.

##### a. Laplace approximation

In the Laplace approximation, the variational distribution is taken to be a Gaussian centered at the maximum a posteriori (MAP) estimate, 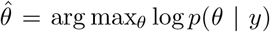, with covariance given by the inverse Hessian of the negative log-posterior at the optimum. The corresponding evidence lower bound (ELBO) is approximated as

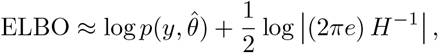

where 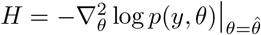 is the Hessian evaluated at the MAP estimate.

##### b. Origin detection

Candidate origin positions are identified from local maxima in the derivative of the smoothed RFD profile (Appendix E). The threshold on the derivative controls how many origins are detected, and the pipeline is run across a range of thresholds to produce candidate origin sets of varying sizes, from which the optimal number is selected via ELBO comparison.

Two peak selection strategies were evaluated: one based on peak height and one based on peak prominence. In the synthetic datasets, where origins are well-separated by construction, both strategies perform similarly, as the dominant peaks are unambiguous. In experimental data, however, closely spaced origins can produce overlapping derivative peaks, and prominence-based selection tends to preferentially retain the dominant peak of a cluster while suppressing its neighbors. Peak height-based selection, by contrast, is more sensitive to closely spaced origins and yielded higher ELBO values on experimental data; it was therefore adopted for the experimental analysis.

##### c. Pruning

Even with height-based detection, once the best model and origin number have been selected via ELBO comparison, the origin set may still contain redundant origins. These are origins with very low observed efficiency *E*_*i*_ that contribute little to the RFD profile but inflate model complexity. To address this, we introduced a pruning step applied to the selected model. Origins are randomly removed with probability proportional to 1 − *E*_*i*_, so that weakly firing origins are preferentially pruned. After each removal, the model is refitted and the new ELBO is compared to the previous one: if the ELBO decreases, the removal is rejected and the previous configuration is restored. This procedure is repeated for a number of rounds equal to the initial number of detected origins. Note that pruning is applied only to experimental data; in the synthetic setting, the known origin positions and well-separated spacing make it unnecessary.

##### d. Full pipeline

The complete analysis pipeline can be summarised as follows: given RFD data, (i) detect candidate origin positions across a range of derivative thresholds (Appendix E); (ii) for each candidate origin set, fit the Exponential and Weibull models with and without delay using the Laplace method; (iii) select the model and origin number that maximise the ELBO; (iv) for experimental data, apply the pruning procedure to the selected model: origins are randomly removed with probability proportional to 1 − *E*_*i*_, the model is re-fitted after each removal, and the removal is accepted only if the ELBO does not decrease; this is repeated for a number of rounds equal to the initial number of detected origins; (v) report the posterior mean and standard deviation of *λ*_*i*_ and 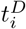 for the final origin set.

Further details on the statistical implementation, including prior distributions and likelihood construction, are provided in Appendix D.

#### 2. Analysis of the performance on synthetic datasets

We evaluated the model selection procedure by simulating data using one model and computing the ELBO for four models: exponential with or without time delay and Weibull with or without time delay. To evaluate model selection over a biological realistic parameter range, we fitted chromosome I of yeast (11 origins) on the experimental data using an exponential and Weibull model with or without delay. The parameters obtained from these fits were used as references for simulations. We then simulated RFD data with a Weibull model with delay (2 A,E) or without time delay (2 B,F) or an exponential model with delay (2 C,G) or without time delay (2 D,H). Noise was added at two different levels, 0.2 and 0.05 RFD units, 0.05 corresponding to our estimation of the experimentally observed noise. We varied the threshold on the derivative of the RFD to select different numbers of origins (ranging from 6 to 24). Laplace method was then used to fit the parameters and estimate the ELBO (Fig. 2). At low RFD noise (0.05) (Fig. 2 A,B,C,D) both the model and number of origin were correctly estimated as they corresponded to the highest ELBO. Only for the simulation with Weibull model the estimated number of origin was lower than the actual number (Fig. 2 B). We found that for the parameters of this model, there where 3 origin with observed efficiency lower than the noise level, which could explain why the estimated number of origin was lower than the actual number. For a number of origins lower than the actual number, the ELBO is very low, maximum around the correct number of origins, and then decreases slowly for a higher number of origins.

**FIG. 2.**
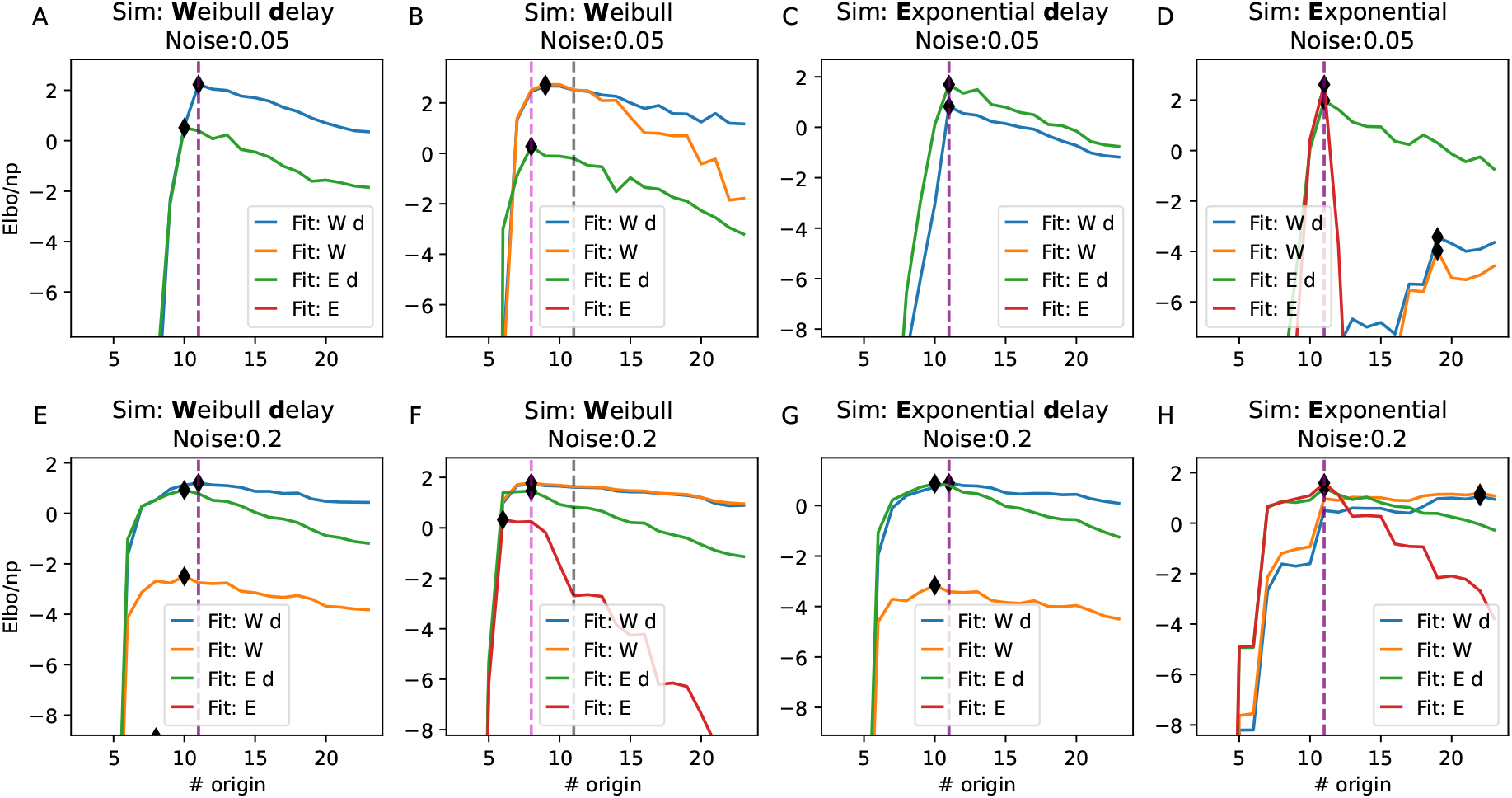
Evaluation of ELBO as a function of the number of origins under different noise levels and firing models. The eight plots are grouped into two panels: A-D represent simulations with low noise 0.05 (in RFD unit), and E-H represent simulations with high noise (0.2). Each subfigure corresponds to data generated with simulation with specific firing model (exponential or Weibull) with or without delay. The curves displayed corresponds to fit with the different models. They include: “Weibull with delay” (blue), “Weibull” (orange), “exponential with delay” (green), “exponential” (red). In some case curves do not appear because their ELBO values are lower than the scale displayed. The black vertical dashed line indicate the total number of origin. The purple one the number of origin with observed efficiency higher than 0.05

For simulations with high noise (Fig. 2 E,F,G,H), the number of origin where most of the time either correctly estimated or underestimated. For simulation with delay (Fig. 2 E,G), both Weibull with delay and Exponential model with delay had similar ELBO. However, model without delay had a much lower ELBO. For simulation without delay (Fig. 2 F,H), the correct model was identified, however, both simulations with or without delay had similar ELBO. As we will see later, the estimated values of the delay parameters were close to 0. Thus, this was not a significant limitation in practice.

We performed this analysis 10 times with random seed. For low noise every time the correct model was inferred and the mean error on the origin number was at maximum of 1.2 origins (SI I). For higher noise, all model but Exponential with delay where correctly identified each time, and the maximum average error was of 2.8 origins.

Overall, the ELBO reliably indicated the firing type and the number of origins, though it was less effective in selecting model without delay when fitting data without delay as models with delay had similar ELBO values.

Now we investigate if the parameters estimated correspond to the ones used for the simulations. As the firing type where correctly identified in a noise level corresponding to experiment, we only focused on correct firing type and possible mismatch for selection of delay. When fitting Weibull simulations with delay using a model with delay (Fig. 3 A), the fit quality improved, the lower the noise. At the experimental noise level all but one origin strength were correctly identified. Only low noise level allowed correct estimation of time delay. For high noise the model sometime overestimates the intrinsic timing and compensate by underestimating the delay.

**FIG. 3.**
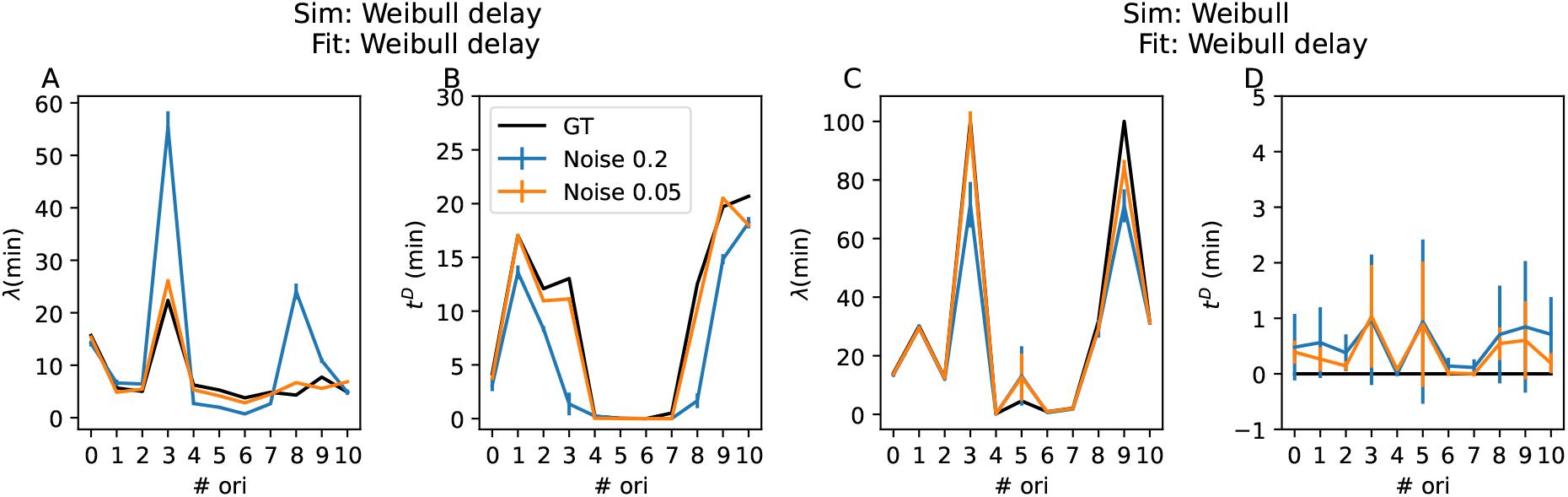
Parameter fitting results for Exponential and Weibull simulations under various noise levels. Fit using the Weibull model with delay for simulation with delay, of origin timing (A) and delay (B). Fit using the Weibull model with delay for simulation without delay, of origin timing (C) and delay (D). In each panel, the blue and orange curves represent fits for Gaussian noise levels of 0.2, and 0.05, respectively. The black line represents the ground truth. The ELBO has been divided by the number of data points fitted.

For Weibull simulations without delay, fitting with a model with delay, provided correct strength and produced accurate parameter estimates (Fig. 3 D). The model correctly predicted a very low delay for all origins.

In conclusion, fitting with the correct model yielded strong parameter estimates. Even with mismatched models, reasonable estimates were obtained, especially when fitting with models containing additional parameters.

#### 3. Fits of experimental data

##### a. Fits of experimental data

To validate our model on real data and determine the number of origins required to reproduce experimental RFD profiles, we applied the full pipeline to the dataset of [12]. Varying the derivative threshold across a range of values revealed a characteristic pattern in the ELBO as a function of origin number: an initial steep increase as genuine origins are added, followed by a slow decrease, and then a second sharp increase driven by the inclusion of redundant origins. This non-monotonic behaviour motivated the use of peak height-based detection combined with the post-selection pruning procedure described above, which resolved the redundancy and identified a clean optimal origin number. The best model across all chromosomes was consistently the Weibull distribution with activation delays, selected on the basis of ELBO comparison. The highest ELBO was obtained for a total of 776 origins.

We illustrate the fit of the data with the best model, using the average of the intrinsic strength and delay, on the first 7 chromosomes (Fig. 4 A). When comparing experimental data with simulated ones, the Pearson correlation ranged between 0.96 and 0.99 with an average correlation of 0.98. The fits for all chromosomes are displayed in Fig 6.

**FIG. 4.**
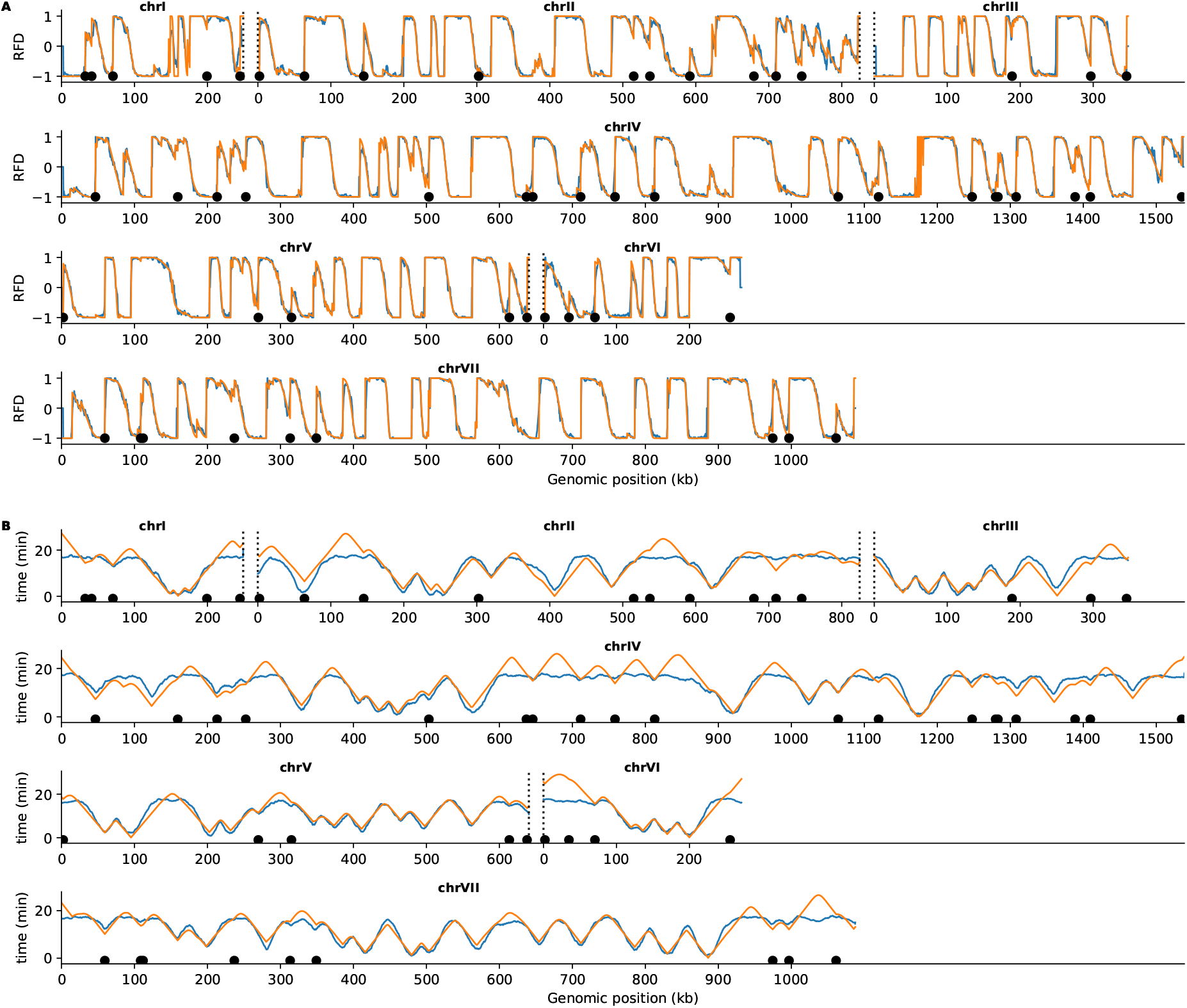
(A) Comparison between experimental (blue) and simulated (orange) RFD, for the first 7 chromosomes. The black dots are highly delayed origins. B) Comparison between simulated MRT (orange) and rescaled Brdu profile (blue).

To further validate the model, we compared the MRT profiles predicted by our framework against an independent experimental dataset. Crucially, the model parameters (*λ*_*i*_ and 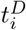) were estimated solely from the RFD data of [12], without any fitting to or knowledge of the BrdU profiles. The predicted MRT was then compared to rescaled BrdU incorporation profiles obtained from nanotiming experiments [24], which provide an independent measure of replication timing. We performed a simple rescaling of the BrdU profiles for display purposes only: the BrdU profiles were standardized giving sBrdU, and we display 20(1 − sBrdU) to match the MRT scale. The agreement between the two is excellent, with Pearson correlation ranging from 0.82 to 0.97 across chromosomes (average 0.90), demonstrating that the model generalizes beyond the RFD data it was fitted to. Some discrepancy arises near telomeres, where the BrdU profiles are flattened, likely reflecting reduced sensitivity of the nanotiming experiment in late-replicating regions rather than a failure of the model.

When looking at the distribution of origin intrinsic strength and delay (Fig. 5 A), we notice that only a small number of origins 145/776 are delayed by more than 3 minute. We choose this as a threshold with respect to our analysis of simulated data, where the estimation of delay for origin without delay never exceeded 3 min, accounting for the average estimate plus one standard deviation. These highly delayed origins have a rather high observed efficiency (Fig. 5 B-C) and have an intermediary replication time (Fig. 5 D). Part of them are located at telomeres (eg chrI right telomere, chrII left telomere, chrIII right telomere).

**FIG. 5.**
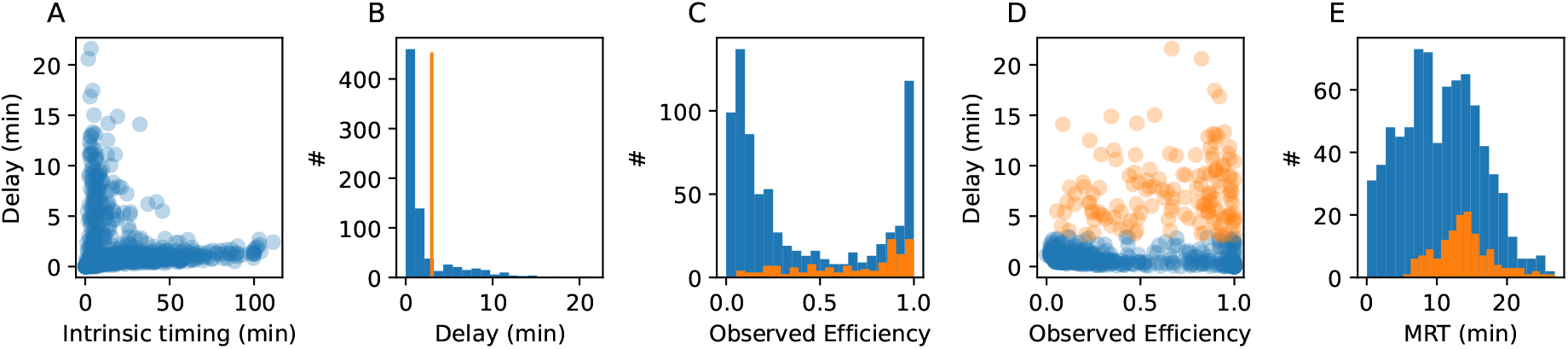
A) Scatter plot of Intrinsic strength vs Delay for the 776 origin selected B) histogram of the delays (blue) vertical line at 3 minutes (orange); C) histogram of the Observed Efficiency for all the origins (blue) and for the 145 origin with a delay higher than 3 minutes (orange). D) Scatter plot of Observed Efficiency vs Delay for all the orgins (blue) and for the origin with a delay higher than 3 minutes (orange). E) Histogram of the replication time of all (blue) and high delay origins (orange)

**FIG. 6.**
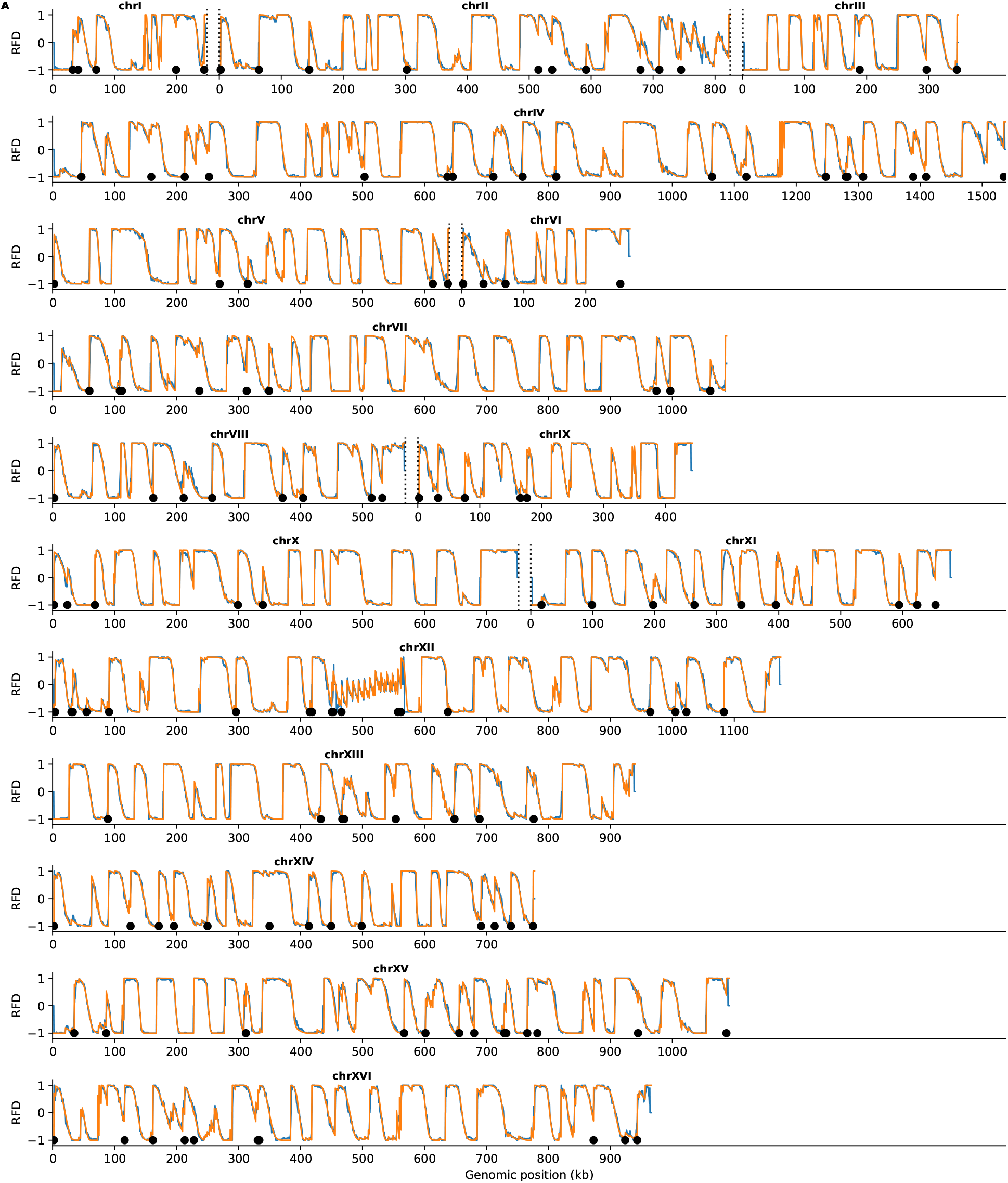
(A) Comparison between experimental (blue) and simulated (orange) RFD, for all the chromosomes. The black corresponds to origins with high delay.

**FIG. 7.**
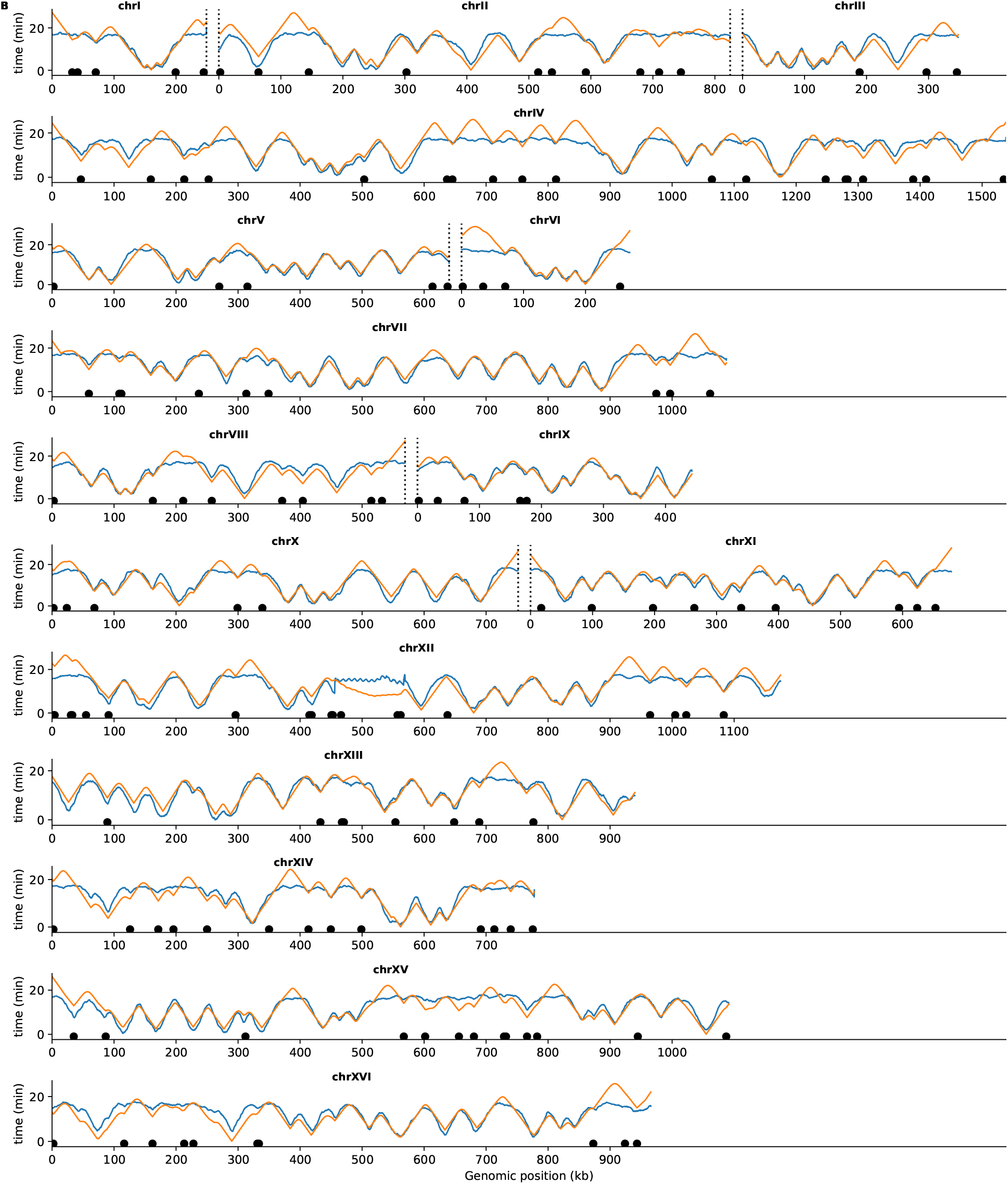
(A) Comparison between experimental Brdu rescaled MRT (blue) and simulated MRT (orange), for all the chromosomes. The black corresponds to origins with high delay.

For those, we think it could be due to the fact that our simulation do not include capture and release of firing factor: we tried fitting our previous model [16] that include recycling of firing factor. For all models the fit were quite good. The best model through ELBO selection was always the Weibull model. The only part that where not almost perfectly fit where located at chromosomes extremities. However, fit with delay were better at chromosomes extremities.

##### b. Fit with only know origins, Comparison origins / detected peaks

We also fitted the chromosomes using Confirmed and Likely origins from the ori db database. The dubious origin where not used because typically their localization is not precise. The best average correlation was of 0.86 so around 0.1 point lower than the best model, clearly due to missing origins, and was obtained for model with delay. Both Weibull and exponential model had similar correlations.

## V. DISCUSSION

### a. Model and its Validation

In this study, we developed analytical models to compute MRT and RFD for genome replication systems with multiple origins, under both exponential and Weibull-distributed firing times. By deriving closed-form expressions for MRT and RFD, we established a computationally efficient Bayesian framework to estimate intrinsic origin timing from experimental data.

Our method yields high correlations (0.96–0.99) between simulated and experimental RFD profiles across yeast chromosomes (Fig. 4), demonstrating its utility for genome-wide replication timing analyzes. Notably, the Weibull model (*k* = 2), which avoids the abrupt firing onset of the exponential distribution, provided marginally better fits to experimental data, as evidenced by higher ELBO values. This aligns with biological expectations, as origin activation in vivo is unlikely to follow a very abrupt start due to the diffusion of factors in the nucleus.

Our Bayesian model selection revealed that a model including activation delays provides a better fit for the experimental data, pointing to a biologically significant subset of origins whose firing is temporally regulated. The remarkable correlation between predicted and experimental RFD profiles, together with the independent validation against BrdU nanotiming data [24], for which no additional fitting was performed, demonstrates the model’s capacity to capture essential features of the replication landscape.

The systematic evaluation of model performance across varying origin detection thresholds provides robust insights into the replication dynamics of *S. cerevisiae*. Our analysis identified ~ 780 origins as optimal for reproducing experimental RFD profiles, consistent with experimental estimates that only 500–600 origins show significant firing under normal conditions [10], probably due to the presence of limiting factors required to activate origins [25]. Our model’s optimal origin number matches these experimental counts, suggesting the estimated *λ*_*i*_ values reflect biological licensing potential filtered through factor availability [26].

### b. Biological Implications

Our distinction between *λ*_*i*_ (intrinsic mean firing time) and *E*_*i*_ (passivation-adjusted firing probability) provides mechanistic insights into how genomic context shapes replication outcomes. While *E*_*i*_ reflects an origin’s net contribution to replication, *λ*_*i*_ represents its inherent activation timescale — a critical parameter for studying origin regulation that is not directly accessible from population-level observables alone.

The relationship *E*_*i*_ = MRT(*x*_*i*_)*/λ*_*i*_ reveals that two origins with identical intrinsic timing *λ*_*i*_ can exhibit markedly different observed efficiencies depending on their genomic neighborhood. An origin embedded among strong, early-firing neighbors will have a low MRT(*x*_*i*_) — it is frequently replicated passively before it has a chance to fire — and will therefore display a low *E*_*i*_ despite its intrinsic potential. Conversely, an isolated origin or one in a late-replicating region will retain a high *E*_*i*_ close to 1. This positional modulation of efficiency, encoded in MRT(*x*_*i*_), explains how identical molecular machinery at two loci can give rise to very different population-level replication behaviors, and underscores why disentangling *λ*_*i*_ from *E*_*i*_ is essential for understanding origin regulation.

Our *λ*_*i*_ parameter directly corresponds to an origin’s ability to recruit limiting initiation factors. This property is determined by ACS motif strength and nucleosome positioning [26, 27] that can be further modulated by chromatin context such as regulation by Rif1 [24, 28], Forkhead transcription factors [29], Dbf4 at the kinetochore [30], or trans elements in a larger 3D chromatin context [29]. Future work should focus on elucidating the molecular mechanisms governing these delays, potentially involving chromatin remodelling, transcription factor binding, or higher-order chromosome structure.

We identified approximately 150 origins with activation delays exceeding 3 minutes. For the subset of these are located at telomeres, their interpretation is less straightforward. Indeed, as discussed in the Limitations section (next paragraph), when firing factor recycling is included in the model, a model with activation delays provides better fits specifically in subtelomeric regions. It is thus possible that the delays inferred there reflect an unmodelled recycling effect rather than genuine regulatory kinetics. The delayed origins located at internal chromosomal positions are not subject to this ambiguity and represent stronger candidates for genuinely regulated activation kinetics.

### c. Limitations and Future Directions

One important hypothesis is that all origins are licensed in each cell, which is not fully consistent with current experimental knowledge. However, the excellent quality of our fits suggests this simplification does not critically affect the results. We simulated synthetic datasets with partial licensing at 80% for either a strong or a weak origin (Fig. 8). From visual inspection, partial licensing of a weak origin produces a signature very similar to that of a fully licensed but rarely activated origin, making the two scenarios indistinguishable from RFD data alone. For a strong origin, however, partial licensing produces a distinctive plateau around − 0.5 in the RFD profile, a signature not observed in our experimental dataset. Our model therefore predicts that, if partial licensing is present, it predominantly affects weak origins. We also proposed a perturbative mathematical framework to incorporate partial licensing as a correction to the full MRT.

**FIG. 8.**
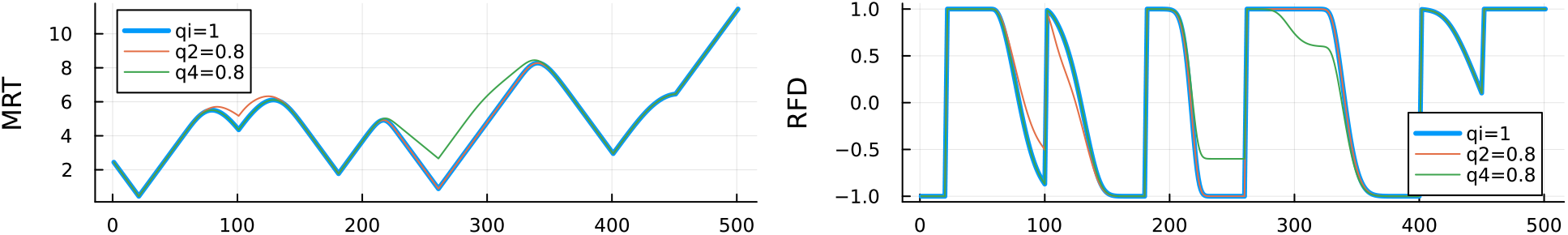
Comparison of a system with full licensing (blue), partial licensing of the second origin (weak), or partial licensing of the fourth origin (strong) for MRT (left) and RFD (right)

While our current framework assumes a constant fork speed, it could be extended to accommodate variable fork speeds through an iterative approach. First performing a fit at constant speed to obtain an initial MRT, then estimating a position-dependent speed from that MRT, and iterating until convergence.

Another limitation is the omission of firing factor recycling, which was shown to be critical for reproducing the temporal firing rate of origins [31]. We investigated this by fitting a model incorporating this feature [16], and found that it primarily affected subtelomeric regions, where a model with activation delays provided better fits. However, at the level of global model selection via ELBO, the model without delays was still preferred, suggesting that the activation delays inferred from experimental data at non-subtelomeric origins are not explained by the omission of firing factor recycling.

Another simplification lies in the variational inference framework, which assumes independence between origin parameters (mean-field approximation). While this enables scalable parameter estimation for large systems (e.g., yeast chromosomes with ~50–100 origins), it neglects potential coupling between neighbor origins, that could compensate each other, and would thus lead to a higher uncertainty on the estimates. The mean-field approximation also impacts model selection via the ELBO, as it underestimates posterior uncertainties. In synthetic tests, the ELBO reliably identified the correct firing type but was less discriminative between models with and without delay. In practice this was not a significant issue, as models fitted with delay on data generated without delay consistently returned delay values close to zero (Fig. 3). Future work could incorporate structured variational distributions modelling pairwise correlations, spatial priors such as Gaussian processes with chromosomal-distance-dependent kernels, or hybrid approaches combining variational inference with Hamiltonian Monte Carlo for small chromosomal regions.

### d. Broader Impact

Our analytical derivations bridge a critical gap between replication theory and data-driven modelling. By eliminating the need for stochastic simulations, we enable Bayesian inference on timescales compatible with high-throughput sequencing datasets. This opens avenues for studying replication dynamics under varying cellular conditions (e.g., stress, cell cycle perturbations) or across species with divergent origin licensing strategies. As single-molecule and sequencing technologies continue to refine spatiotemporal replication maps, integrating biophysical correlations into Bayesian frameworks will be essential to unravel the complexity of origin regulation in higher eukaryotes.

## Appendix A: Deriving MRT from *p*_*i*_

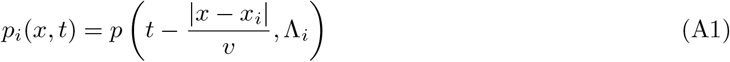

The probability that another fork *j* arrives at *x* later than *t* is given by

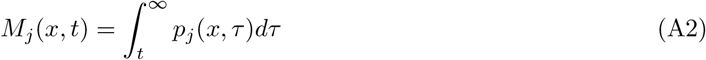

So the probability that the fork coming from *i* replicate *x* at *t* is

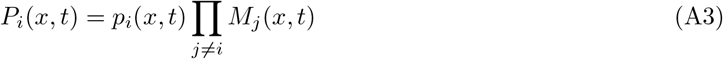

And the probability for a point *x* to be replicated at *t* is

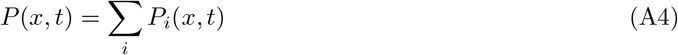

The latter definition forms the base of Retkute al [15] model. Noticing that 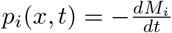 allows to write *P* (*x, t*) with a denser form:

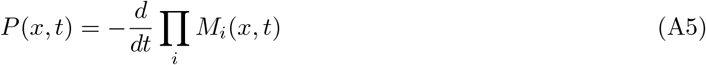

In fact *s*(*x, t*) = Π *_i_ M*_*i*_(*x, t*) is the probability that x has not been replicated at time *t*. This relationship as already noted in [32]. Eq. (A5) allows us to write a simplified version of the MRT. The MRT is defined by:

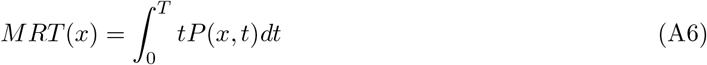

with *T* a time that can be taken large with respect to the S-phase but that we will push to infinity at the end of this derivation. Using Eq. (A5):

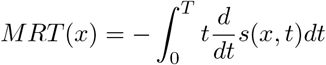

and if we integrate by parts and using *s*(0, *x*) = 1 because no DNA as been replicated at time *t* = 0.

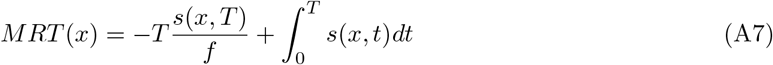

As *s*(∞, *x*) tends towards zero faster than *t* because it is the product of several cumulative distributions, this simplifies Eq. (A8) :

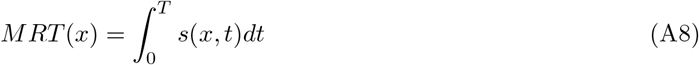

## Appendix B: Exponential case

If we consider that the origins follow an exponential law of firing, starting at 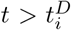, then 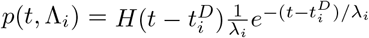 and the cumulative distribution is given by 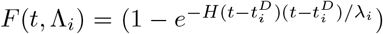 with H(t) the Heaviside function.

And so

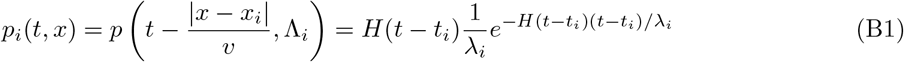

and

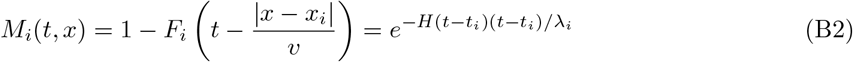

with 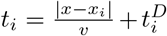 the total delay containing two factors. 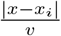 corresponding to the time for the fork to propagate and 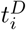 is an optional delay that can be added to mimic a delay in activation of the origin.

To determine *MRT* (*x*), we class the *n* origins by increasing time of arrival to x, *t*_*i*_. So 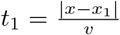 is the closest origin to x. And we cut the integration into pieces: from (0 to *t*_1_), all *M*_*i*_(*t, x*) are equal to 1. Between *t*_1_ and *t*_2_ only *M*_1_(*t, x*) is different from one and: 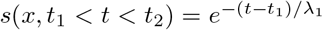. And more generally: 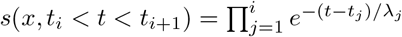. We also define *t*_*n*_ = ∞, thus

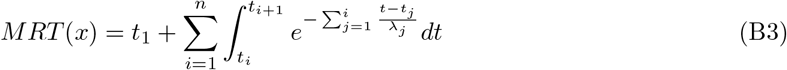

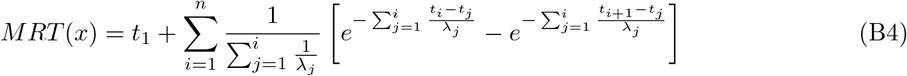

if we define with 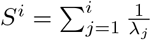

then one can show that 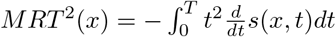 is equal to:

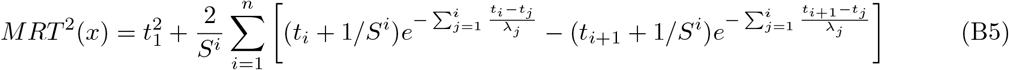

so that the dispersion of the MRT around its mean value is easily computed when computing the MRT.

To derive *E*_*i*_ the observed efficiency, we can notice that given the exponential form of the firing law, 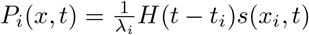. As 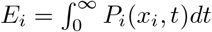 then

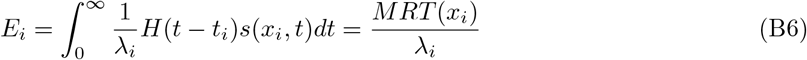

This is true in the case where there is no delay of activation 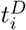 because then *t*_*i*_ = 0 as the origin is located at position *x*_*i*_. This relationship that link the observed efficiency, the intrinsic strength *λ*_*i*_ was derived using several approximations in Arbona et al. [16] using a more complex model, and is here a consequence of the exponential firing of the origin.

In the case where 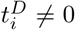, if the index of *t*_*i*_ is k. if k is different from zero, it means that the origin i is activated after forks from origins arrive first at *x*_*i*_ then:

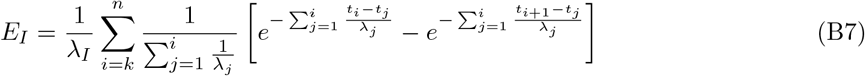

RFD is defined by the derivative of the MRT with respect to *x*: 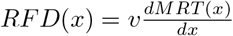. In Eq. B4 the dependency as a function of *x* is implicit in the *t*_*i*_ terms. As *t*_*i*_ = |*x*−*x*_*i*_|*/v* and 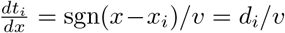with d stating for direction

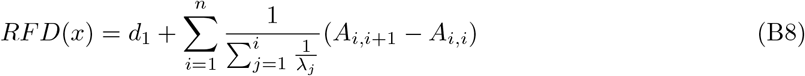

with

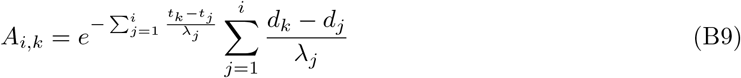

## Appendix C: Analytical formulas Weibull distribution with *k* = 2

For the Weibull distribution with *k* = 2, the probability for an origin to fire 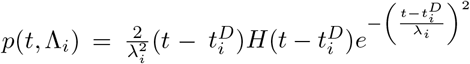 and the cumulative distribution is 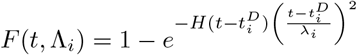.

By sorting again the origin by time of arrival:

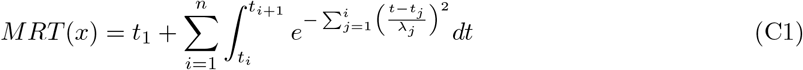

and by completing the square of the Gaussian integral one get

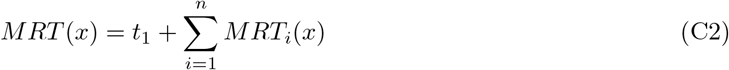

with

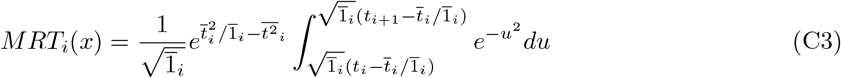

with 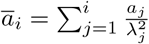 for any list *a* = (*a*, *a*, …, *a*_*N*_) and 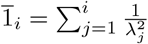. The integral part of *MRT* (*x*) is the Gauss error function between two times which can be computed quickly as good analytical analytical approximation are known.

For the second moment:

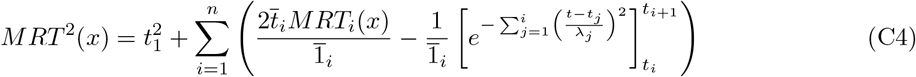

For the Observed efficiency unfortunately the relationship with *MRT* (*x*) is not as simple as in the exponential case. First the *MRT*_*i*_(*x*) term can be decomposed in *MRT*_*i*_(*x*) = *F*_*i*_*I*_*i*_ with

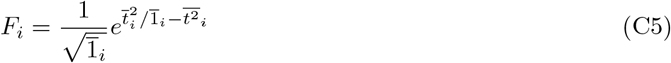

and

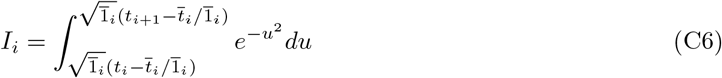

So that the efficiency of origin *l* is:

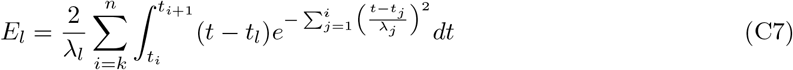

where the sum starts at *k*, the index of *t*_*l*_, as in the exponential case.

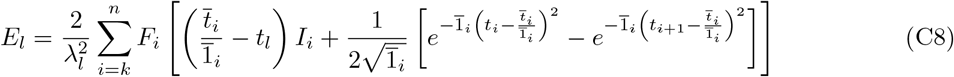

For the RFD,

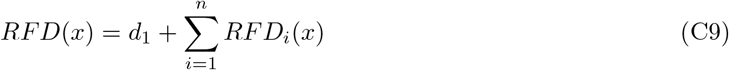

with

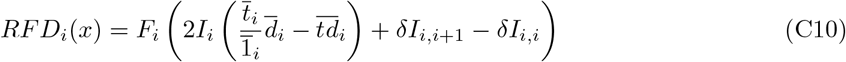

with

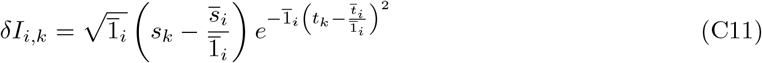

## Appendix D: Detail of parameters prior and likelihood for the RFD

In practice we perform 10 optimisations with random seed and select the optimisation result with higher ELBO. Priors to the parameters *λ*_*i*_ were chosen as Weibull (k=2) efficiencies (mean = *T*_*s*_*/*2) and exponential-distributed activation delays with mean = *T*_*s*_*/*5 for models with delay. Typically RFD residuals were modeled as 𝒩(0, *σ*). Also we noticed that stabler result where obtained when fitting not the RFD but 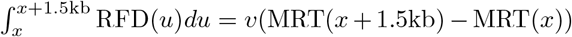. As a consequence on synthetic data, where we simulated data at a 100 bp resolution and then added some gaussian noise 𝒩(0, noise) the error *σ* was estimated as *noise/nbins*^0.5^, where noise is the value of the introduced noise and *nbins* represents the number of bins in 1.5 kb. This is because for this simulated data, the noise at each bin is uncorrelated. For experimental data, the error for each bin was estimated as *p*(*x*)(1 − *p*(*x*))*/*(*n*(*x*) + 20), where *p*(*x*) = (*RFD*(*x*) + 1)*/*2 and *n*(*x*) the number of fork in bin *x* (20 was added to the fork number to prevent overestimation of the confidence interval). This quantity *p*(*x*) can be seen as the probability for a fork to go to the right. However, contrary to the simulated data we used *σ* = *p*(*x*)(1 − *p*(*x*))*/*(*n*(*x*) +20) without dividing by the square of the number of bins, as the data coming from single cell molecule is highly correlated on the 1.5 kb scale.

## Appendix E: Origin Detection Algorithm

Origin positions were identified through peak detection on the replication fork directionality (RFD) derivative. The pipeline comprises three steps:

1. **Smoothing**: RFD profiles were convolved with a Gaussian kernel (*σ* = 3 kb) to reduce high-frequency noise:

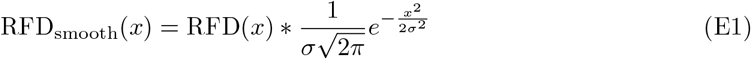
2. **Differentiation**: The numerical derivative was computed using central differences:

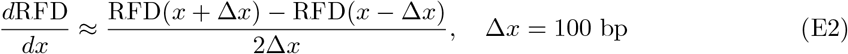
3. **Peak Detection**: Local maxima in |*d*RFD*/dx*| were identified using SciPy’s find peaks with:
  - Minimum prominence: 0.001 per 100 bp
  - Minimum inter-peak distance: 3 kb
  - Height threshold: *>* 5*×* median absolute deviation

## Appendix F: Fit all

**Table I.** Model performance and origin error statistics under different noise levels.

| Noise | Data | Correct | Mean | Median | Exact | Max |
| --- | --- | --- | --- | --- | --- | --- |
| 0.2 | Weibull (Delay) | 10/10 | -0.10 | 0.0 | 7/10 | -1 |
| 0.2 | Weibull | 10/10 | -2.80 | -3.0 | 0/10 | -4 |
| 0.2 | Exponential (Delay) | 3/10 | -0.30 | 0.0 | 7/10 | -1 |
| 0.2 | Exponential | 10/10 | -0.10 | 0.0 | 9/10 | -1 |
| 0.05 | Weibull (Delay) | 10/10 | 0.00 | 0.0 | 10/10 | 0 |
| 0.05 | Weibull | 10/10 | -0.80 | -1.0 | 2/10 | -2 |
| 0.05 | Exponential (Delay) | 10/10 | 0.10 | 0.0 | 9/10 | 1 |
| 0.05 | Exponential | 10/10 | 0.00 | 0.0 | 10/10 | 0 |

